# Cap-related modifications of RNA regulate binding to IFIT proteins

**DOI:** 10.1101/2024.06.25.600627

**Authors:** Jingping Geng, Magdalena Chrabaszczewska, Karol Kurpiejewski, Anna Stankiewicz-Drogon, Marzena Jankowska-Anyszka, Edward Darzynkiewicz, Renata Grzela

**Author notes:** **Correspondence to Renata Grzela:** Division of Biophysics, Institute of Experimental Physics, Faculty of Physics, University of Warsaw, Pasteura 5, 02-093 Warsaw, Poland. These two authors contributed equally to this work (Jingping Geng, Magdalena Chrabaszczewska).

## Abstract

All cells in our body are equipped with receptors to recognize pathogens and trigger a rapid defense response. As a result, foreign molecules are blocked and cells are alerted to the danger. Among the many molecules produced in response to viral infection are interferon-induced proteins with tetratricopeptide repeats (IFITs). Their role is to recognize foreign mRNA and eliminate it from the translational pool of transcripts. In the present study, we used biophysical methods to characterize the interactions between IFIT1 protein and its partners IFIT2 and IFIT3. IFIT1 interacts with IFIT3 with nanomolar binding affinity, which did not change significantly in the presence of the preformed IFIT2/3 complex. The interactions between IFIT2 and IFIT3 and IFIT1 and IFIT2 were one order of magnitude weaker. We also present kinetic data of the interactions between the IFIT protein complex and short RNA bearing various modifications at the 5’ end. We show kinetic parameters for interaction between IFIT complex and RNA with m^6^A_m_ modification. The results show that the cap adjacent m^6^A_m_ modification is a stronger signature than cap1 alone. It blocks the formation of a complex between IFIT proteins and m^7^Gpppm^6^A_m_-RNA much more effectively than other cap modifications. In contrast, m^6^A in the 5’UTR is not recognized by IFIT proteins and does not contribute to translation repression by IFIT proteins. The data obtained are important for understanding the regulation of expression of genetic information. They indicate that 2’-*O* and m^6^A_m_ modifications modulate the availability of mRNA molecules for proteins of innate immune response.

## Introduction

Innate immunity is an organism’s first line of defense against pathogens. Pathogen-associated molecular patterns (PAMPs) are recognized by specific pattern recognition receptors (PRRs), which are activated upon binding of the PAMP and induce various pathogen-specific responses (Agrawal et al. 2003; Hornung and Latz 2010). Among effective PAMPs are foreign nucleic acids characterized by the presence of dsRNA structures or the absence of specific host modifications (Nagaraj et al. 2022). The signature of ‘self’ RNA is the cap structure at the 5ʹ end, consisting of 7-methylguanosine linked to the RNA body by a triphosphate chain, with 2ʹ-*O*-methylation at the first (cap1) or first and second (cap2) nucleotides (Hyde and Diamond 2015; Fensterl and Sen 2015). The 2ʹ-*O*-methylation of the first transcribed nucleotide of cellular mRNA is catalysed by 2′-*O*-methyltransferase 1 (CMTR1), a protein upregulated during viral infection and responsible for the activation of expression of other interferon stimulated genes (ISGs) (Terenzi et al. 2006). The CMTR1 depletion leads to a decrease in the production of ISGs-encoded proteins even though the amount or stability of the transcripts encoding these proteins does not change. Moreover, CMTR1-regulated ISGs transcripts are inhibited by the IFIT1 sensor in cells depleted of CMTR1 (Williams et al. 2020). The presence of 2′-*O*-methylation of the first transcribed nucleotide significantly reduces the stability of the RNA/IFIT1 complex, thus, translation of the modified transcript is not blocked (Miedziak et al. 2019). One of the most common modifications in the RNA chain is *N*6-methylation of adenosine nucleotide (Brocard et al. 2017). It is a dynamic modification, introduced by methyltransferases (writers) (Bujnicki et al. 2002; Yang et al. 2018) and removed by demethylases (erasers) (Zheng et al. 2013; Wei et al. 2018). The identification of m^6^A is mediated by m^6^A-binding proteins (readers) (Xiao et al. 2016; Shi et al. 2017; Du et al. 2016) and proteins that bind mRNA in the absence of m^6^A (anti-readers) (Edupuganti et al. 2017; Li et al. 2018). This modification plays multiple roles, affecting maturation, stability, splicing, export, translation and finally RNA decay (Batista et al. 2014). Additionally, it regulates pathogen-host interactions (Tan and Gao 2018) with the presence or absence of m^6^A influencing viral gene expression during infection (An et al. 2020; Lichinchi et al. 2016; Li et al. 2021; Kennedy et al. 2017).

The adenosine nucleotide can also be methylated to the *N*6,2ʹ-*O*-dimethyladenosine (m^6^A_m_) form. Cap-adjacent m^6^A_m_ (CA-m^6^A_m_) is formed as a result of the activity of PCIF protein (phosphorylated CTD-interacting factor 1, also known as CAPAM, cap-specific adenosine methyltransferase) that adds methyl group to the A residue already bearing the 2ʹ-*O*-methylation (Akichika et al. 2019; Boulias et al. 2019; Sendinc et al. 2019; Sun et al. 2019). There is a legitimate hypothesis that CA-m^6^A_m_ affects the binding strength of proteins that regulate mRNA metabolism (Drazkowska et al. 2022), however, there is a limited amount of research determining which cap-binding proteins are sensitive to the presence of the CA-m^6^A_m_. Notably, the epitranscriptomic pattern of both m^6^A and 2′-*O* modifications in the host plays crucial role in the regulation of both mRNA metabolism and gene expression in response to viral infection (Dang et al. 2019; Williams et al. 2020).

Among the earliest produced proteins responsible for viral RNA recognition and binding are IFIT proteins, encoded by IFN-stimulated genes (ISGs). IFIT1 identifies and binds the 5′ end of foreign RNA leading to direct inhibition of mRNA translation (Abbas et al. 2017). It is known that the strength of binding of the RNA molecule by IFIT proteins is affected by the IFITs assembling into a complex. Although certain IFIT proteins like IFIT5 function independently, IFIT1 forms complexes with IFIT2 and IFIT3 (Fleith et al. 2018; Johnson et al. 2018). IFIT1, IFIT2 and IFIT3 can assemble into a complex with a molecular mass corresponding to a trimer or tetramer (150 - 200 kDa). Interactions within this complex exhibit distinct characteristics, in particular, a strong and specific interaction occurs between IFIT1 and IFIT3, mediated by the YExxL motif located at the C-terminus of IFIT3. This interaction is rapid and temperature-independent. By contrast, the interaction between IFIT1 and IFIT2 is weak, time-dependent and influenced by temperature. Nevertheless, IFIT2 forms a stable heterodimer with IFIT3, which subsequently interacts with IFIT1, resulting in the most stable trimeric complex among these proteins (Fleith et al. 2018). However, biophysical data on the interactions between the proteins of the complex are still missing.

Our previous studies thoroughly characterized the direct kinetic interactions between IFIT1 and IFIT5 proteins and RNAs with various native and synthetic modifications at the 5′ end (Miedziak et al. 2019). Notably, IFIT1’s interaction is strictly dependent on the presence of a triphosphate bridge, crucial for electrostatic interactions with protein side chains and most likely initiates ligand recognition (Miedziak et al. 2019; Abbas et al. 2017). RNAs capped with GpppN and m^7^GpppN (cap0) form highly stable complexes with IFIT1, characterized by a very long residence time of the protein on the RNA ligand. Furthermore, the presence of adenozine at the N1 position stabilizes the RNA/IFIT1 complex.

It is known that IFIT1 in complex with IFIT3 or as an IFIT1/IFIT2/IFIT3 trimer has increased ability to bind cap0 RNA (Fleith et al. 2018; Johnson et al. 2018). In primer elongation inhibition assay the trimer binds cap0-RNA twofold more strongly than IFIT1, however authors stated that this data might be biased by saturation of RNA binding. In turn, experiments in the RRL system have shown that the IFIT1/IFIT2 complex inhibits translation of reporter genes at 1.35 fold lower concentrations than IFIT1 alone (Fleith et al. 2018). The IFIT1/IFIT3 complex also has a higher affinity for pppRNA and cap1-RNA than IFIT1 alone, due to the ability of IFIT3 to induce a tighter conformation of the IFIT1 on RNA (Abbas et al. 2017).

The methods used so far provided only approximate parameters characterizing IFIT-RNA interactions. Therefore, we adopted the interferometry-based approach (Miedziak et al. 2019) to study how IFIT2 and IFIT3 enhance IFIT1 binding to capped RNA. Moreover, there are limited data on the interaction of IFITs with RNA bearing important modifications like CA-m^6^A_m_. We synthesized in vitro RNAs with a series of modifications at the 5’ end bearing A, A_m_ or m^6^A_m_ as the first transcribed nucleotide. We performed biophysical characterization of the interactions of IFIT complexes with these RNAs and compared the results with those obtained for IFIT1 alone. These findings are important for understanding the process of gene expression regulation, since both 2’-*O-*methylation and CA-m^6^A_m_ modification modulate the availability of mRNA for translation factors.

## Results

To study the interactions between RNA and the IFIT complex, we prepared proteins: IFIT1, IFIT2, and IFIT3 and formed them into IFIT heterocomplexes. We examined the stability and interactions between individual components of the complex and found out that temperature and time are crucial for proper assembly. These data allowed us to optimize the conditions for the formation of IFIT complexes and to further assess the ability of these complexes to bind capped mRNA.

### Thermal Stability of IFIT Proteins Measured by Differential Scanning Fluorimetry (DSF)

We measured protein stability at different temperatures using nanoDSF with a dynamic light scattering (DLS) module to detect early aggregate formation. IFIT1 aggregated at 42.20 ± 0.22 °C, IFIT2 at 37.90 ± 0.65 °C, while IFIT3 showed no tendency to aggregate (Figure 1A). IFIT2 had the lowest thermal stability, with both the lowest melting temperature (*T*_m_) and aggregation temperature (*T*_agg_), both around 37 °C (Figure 1A and 1B). For this protein we also observed some fluorescence fluctuations and a small peak in the first derivative curve at around 30 °C (Figure 1B), which indicates a low content of additional oligomeric forms. Based on these results, further measurements for IFIT complex assembly were tested at 4 °C and 15 °C.

**Figure 1.**
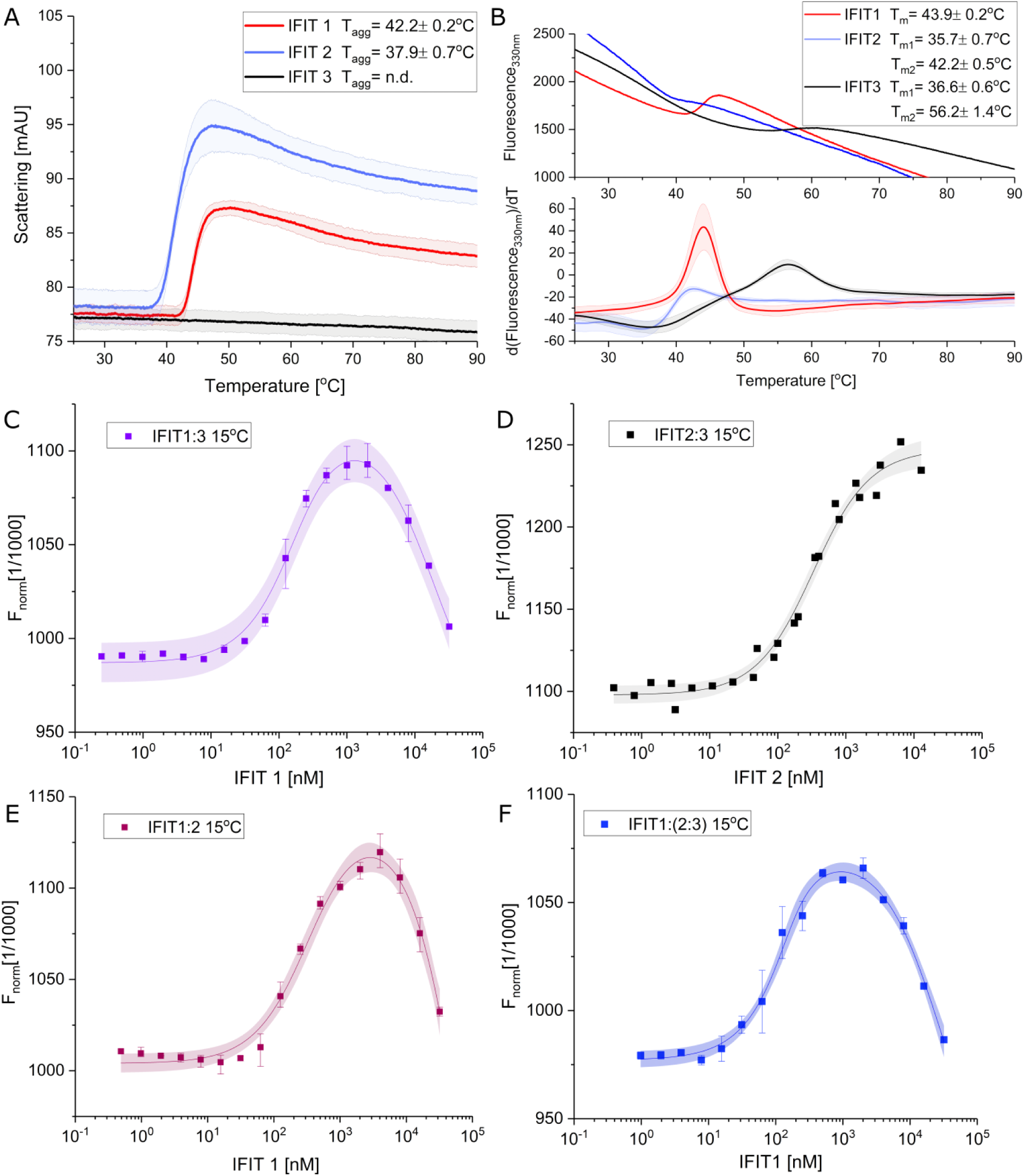
Analysis of IFIT proteins and formation of IFIT complexes. **A.** Aggregation of IFIT proteins monitored by DLS module parallel to the nanoDSF fluorescence measurement. *T*_agg_ – onset temperature of protein aggregation. Results are presented along with the range of standard deviation, shown as semi-transparent color. **B.** Thermal unfolding of IFIT proteins, presented as raw fluorescence intensity at 330 nm (upper panel) and the first derivative (lower panel), both plotted against increasing temperature. *T_m_* calculated for individual IFIT proteins are shown in figure legend. **C-F**. Representative MST pseudo-titration data for the binding of IFIT proteins. Formation of the IFIT complexes measured at 22 °C, after 30 min pre-incubation at 15 °C. Squares represent experimental points, solid lines represent results of fitting and color area represents 95% confidence bands for used model. Titration curves of IFIT3 (**C**), IFIT2 (**E**) and IFIT2/IFIT3 (**F**) with increasing concentrations of IFIT1 ligand. (**D**) Titration curve of IFIT3 with IFIT2 ligand.

### Binding Affinities of IFIT proteins by Microscale Thermophoresis (MST)

An MST assay was used to study the interactions of IFIT1, IFIT2 and IFIT3 proteins. His-tagged IFIT proteins (IFIT2 or IFIT3) were labeled with a fluorescent dye RED-tris-NTA 2^nd^, while the other IFIT protein acted as a ligand. IFIT complexes were assembled under different conditions: without incubation or with prior incubation at 4 °C and 15 °C, the subsequent interaction was measured at 22 °C. Measurement attempts at higher temperatures failed due to adsorption of proteins on the capillary scanner, aggregates on the time response curves, poor reproducibility and lack of stable complexes. The *K*_D_ values for all complexes are summarized in Table 1.

**Table 1.**
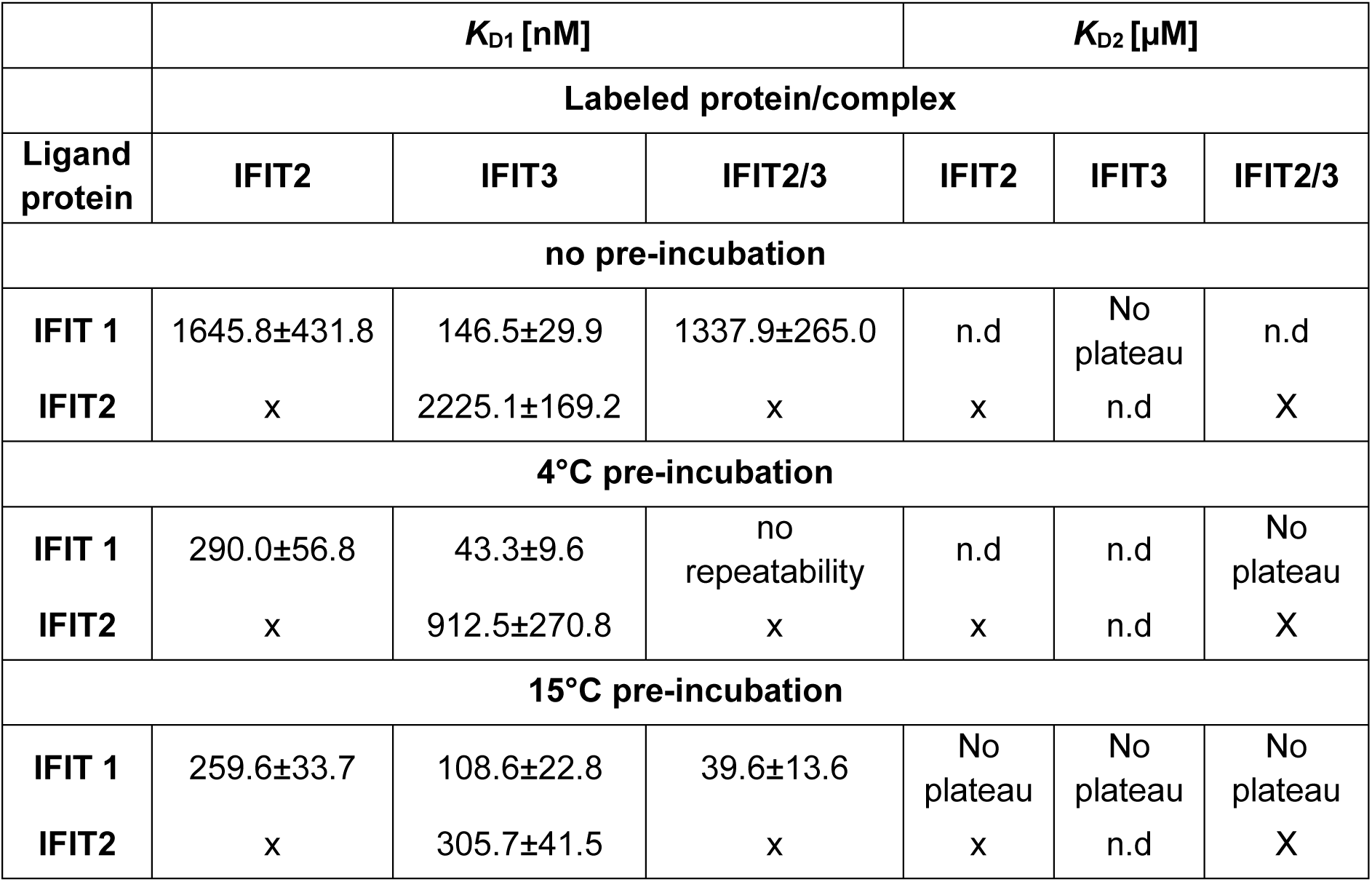
*K*_D1_ and *K*_D2_ binding constants measured by MST for IFIT complexes prepared in different pre-incubation conditions – without pre-incubation and with pre-incubation at 4 °C or 15 °C. All values were determined for measurements at 22 °C; n.d stands for “not detected”.

The strongest binding of IFIT1 and IFIT3 occurred after pre-incubation at 4 °C, with the binding constant *K*_D1_ = 43.3 ± 9.6 nM (Figure S1a). At 15 °C we obtained a non-monotonic binding curve with the first strong binding site, although weaker than at 4 °C (*K*_D1_=108.6 ± 22.8 nM), and a second weak binding site with *K*_D2_ in the micromolar range (Figure 1C). In the experiment without pre-incubation, a monotonic binding curve without a final plateau was observed, with a binding constant *K*_D1_=146.5 ± 29.9 nM (Figure S1a). The results indicate that the IFIT1/IFIT3 complex is assembled very rapidly, is characterized by strong binding and pre-incubation at 4 °C provides the most favorable conditions for the complex formation.

The best conditions for assembly of stable IFIT2/IFIT3 complex were achieved after pre-incubation at 15 °C, with the binding constant *K*_D1_ = 305.7 ± 41.5 nM (Figure 1D). Lower temperature and lack of incubation resulted in weaker binding: *K*_D1_ = 912.5 ± 270.8 nM and 2225.1 ± 169.2 nM, respectively (Figure S1b).

The best conditions for the assembly of IFIT1/IFIT2 complex proved to be pre-incubation at 15 °C, with the binding constant *K*_D1_ = 259.6 ± 33.7 nM (Figure 1E). Similar to the IFIT1/IFIT3 complex, the binding curve showed a second weak binding site with *K*_D2_ in the micromolar range. Pre-incubation at 4 °C resulted in a slightly higher *K*_D1_ = 290.0 ± 56.8 nM, but with greater measurement uncertainty, whereas lack of pre-incubation resulted in very weak binding with *K*_D1_ = 1645.8 ± 431.8 nM (Figure S1c).

The binding affinity of IFIT1 to the heterodimeric IFIT2/IFIT3 complex, preformed under the conditions described above, was measured at 4 °C, 15 °C or without pre-incubation. The strongest binding affinity was observed after pre-incubation at 15 °C. MST measurements showed non-monotonic binding curve with two inflection points, indicating two binding sites: the first very strong with *K*_D1_ = 39.6 ± 13.6 nM and the second much weaker, with *K*_D2_ in the micromolar range (Figure 1F). Measurements after pre-incubation of the IFIT1/IFIT2/IFIT3 complex at 4 °C were not reproducible in duplicate, suggesting that these conditions did not allow the formation of a stable final form of this complex, similar to measurements without pre-incubation (Figure S1d). These results suggest that a certain amount of thermal energy is required for the successful assembly of the complex between IFIT1 and IFIT2/IFIT3.

### BLI Interaction Studies of IFITs with capped mRNA

Using a biolayer interferometry technique, we measured the interactions between a single IFIT1 protein or the IFIT complexes and RNAs with various modifications: cap0 (m^7^GpppAG_RNA_16_), cap1 (m^7^GpppA_m_G_RNA_16_), CA-m^6^A_m_ (m^7^Gppp m^6^A_m_G_RNA_16_) and cap0 plus all A replaced with m^6^A in the RNA body (m^7^GpppAG_m^6^A_RNA_16_).

Control measurements for IFIT1/m^7^GpppAG_RNA_16_ and IFIT1/m^7^GpppA_m_G_RNA_16_ showed slightly higher values than previously published (1.72 - 1.88 times for *K*_D_, 3.77 - 4.12 times for association rate constant (*k*_a_) and 1.96 - 2.18 times for dissociation rate constant (*k*_d_)), (Miedziak et al. 2019), but preserved parameter correlations (Table 2, Figure 2). Around 5-fold weaker *K*_D_ was obtained for m^7^Gpppm^6^A_m_G vs m^7^GpppA_m_G and as much as 32-fold weaker *K*_D_ for m^7^Gpppm^6^A_m_G vs m^7^GpppAG, proving lower affinity of thus modified RNA to IFIT1. For RNA with m^7^GpppAG and body with all A replaced to m^6^A, no significant change in the kinetic parameters of the interaction was observed.

**Figure 2.**
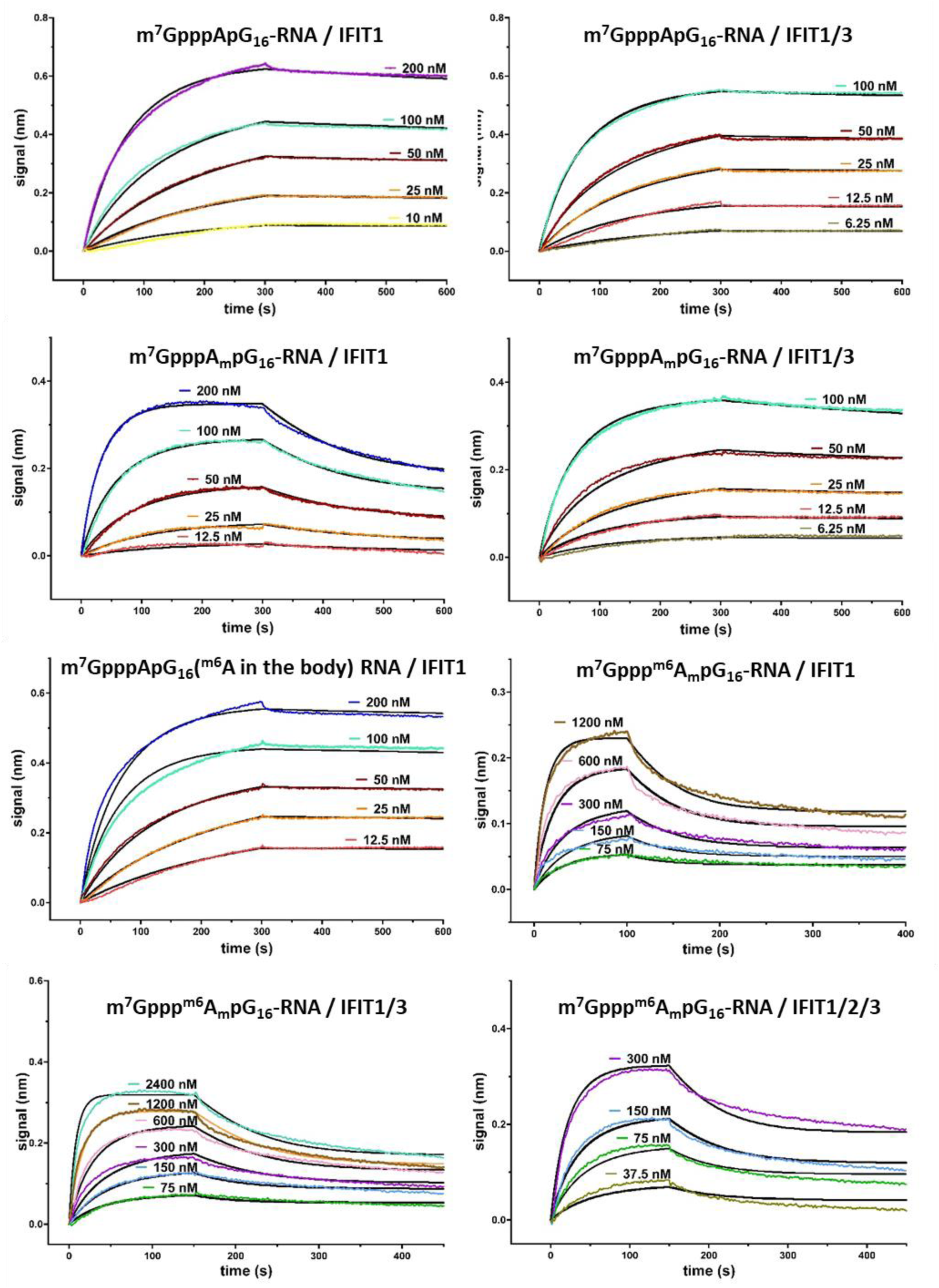
BLI analyses of IFIT proteins interaction with immobilized short RNAs capped with different cap analogues. Biotinylated RNAs bearing cap analogues on 5ʹ end were immobilized on streptavidin sensors and allowed to interact with increasing concentrations of IFIT proteins. The simple 1:1 binding model (black lines) was fitted to BLI data traces (differently colored lines) and plotted as the spectral nanometer shift as a function of time.

**Table 2.**
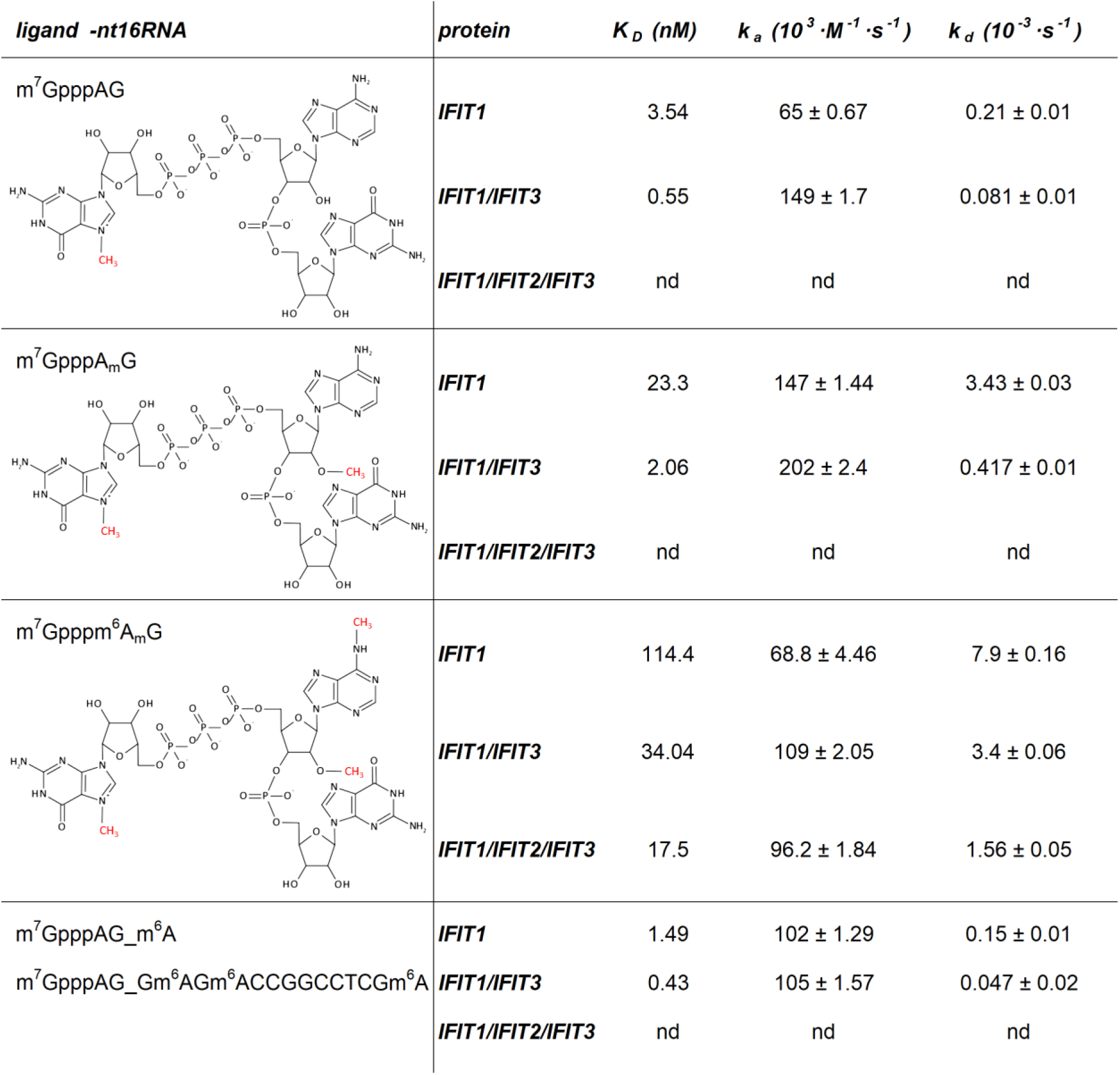
Binding kinetic parameters of the interaction of IFIT1, IFIT1/IFIT3 or IFIT1/IFIT2/IFIT3 with differently capped mRNAs. Displayed *K*_D_, *k*_d_ and *k*_a_ values represent the average of three replicate experiments. nd stands for “not determined”.

RNAs bearing all caps studied showed increased binding affinity to IFIT1 when it was in complex with IFIT3 or IFIT2/IFIT3 (Table 2, Figure 2). Unfortunately, it was not possible to determine the binding constants for IFIT1/IFIT2/IFIT3 complexes with capped RNA (except for m^7^Gpppm^6^A_m_G_RNA_16_), as they were beyond the capabilities of BLI apparatus. By optimizing experimental conditions and understanding the thermal stability and binding affinities of IFIT proteins, we have provided detailed insights into the formation and interactions of IFIT complexes with RNA. Our results proved that the CA-m^6^A_m_ modification provides several times stronger protection against binding of IFIT complexes than cap1 alone and is a very strong signature regulating mRNA translation by IFIT proteins.

## Discussion

Type I interferons, in response to viral infection, induce the expression of ISGs that limit viral infection. IFITs are an example of proteins that are mass produced, and responsible for the inhibition of viral protein synthesis, during the activation of antiviral response (Mears and Sweeney 2018; Fensterl and Sen 2015). IFITs act on two levels, by binding to the eIF3 factor and affecting the general translation in the cell, as well as by specific binding of foreign RNA and consequent prevention of its translation (Vladimer et al. 2014).

Previous reports have shown that IFIT1 acts in complex with IFIT2 and IFIT3 (Mears and Sweeney 2018; Choi et al. 2018; Fleith et al. 2018; Johnson et al. 2018). Therefore, we reconstituted complexes of these proteins for the study of their interactions with RNA. The IFIT1/IFIT3 complex was obtained after pre-incubation at low temperature. In contrast, the formation of the IFIT2/IFIT3 complex required a certain amount of thermal energy. Using MST, we found strong nanomolar interaction between IFIT1 and IFIT3, which did not change significantly when IFIT1 interacted with IFIT2/3 preformed complex. The interactions between IFIT2 and IFIT3 and IFIT1 and IFIT2 were one order of magnitude weaker.

IFITs have the ability to specifically bind foreign RNA to prevent its translation. Determinants of proper recognition of “self” – “non-self” mRNAs are therefore extremely important for the organism’s defense against pathogens. To date, various signatures of foreign mRNA have been identified, for example, the presence of triphosphate at the 5’ end of the RNA or cap0 modification (Daffis et al. 2010; Züst et al. 2011; Abbas et al. 2017). Interestingly, during viral infection, modifications at the 5’ end of host and pathogen transcripts are rearranged (Williams et al. 2020). As a result of the antiviral response, the CMTR1 protein is produced, which catalyzes the 2’-*O*-methylation of the first transcribed nucleotide. The introduction of cap1 modification into transcripts encoding ISG proteins, protects them from recognition by IFIT1 protein. Thus, cap1 modification contributes to the recognition of “self” RNA and to the expression of specific ISGs that further help fight viral infection. It is important to note here that the role of cap1 modification goes beyond marking mRNA as ‘self’ and serves as a mechanism for epitranscriptomic regulation of gene expression. It has been demonstrated that CMTR1-catalyzed methylation of cellular transcripts is essential for promoting histone and ribosomal gene expression in the process of differentiation of embryonic stem cells (Liang et al. 2022) and in the development and fertility of mammalian embryos (Dohnalkova et al. 2023). 2′*O*-methylation of specific ISGs has also been reported to play a critical role in dendritic cell morphogenesis and brain development (Lee et al. 2020). Thus, this modification has wide-ranging cellular significance for both proper RNA recognition and finely tuned gene expression. Other studies have reported that only the presence of both cap1 and cap2 modifications provide effective protection against binding by IFIT proteins (Abbas et al. 2017). Recently it was reported that mRNA with cap2 or m^6^A_m_ at the 5ʹ end did not co-precipitate with IFIT1 protein when transfected into A549 cells (Drazkowska et al. 2022), suggesting no recognition of cap adjacent m^6^A_m_ (CA-m^6^A_m_) modification by IFIT1.

m^6^A_m_ affects processes such as RNA degradation and translation. Depending on the type of experiment, the cell line used or the type of transcript, positive or negative effects of CA-m^6^A_m_ modification on translation and mRNA stability have been observed (Mauer et al. 2017; Sendinc et al. 2019; Sikorski et al. 2020). First, it was shown that m^6^A_m_ promotes mRNA stability (Mauer et al. 2017) and increases expression levels (Linder et al. 2015; Akichika et al. 2019). Further studies confirmed these conclusions in JAWSII cells, while studies in 3T3-L1 and HeLa showed conflicting effects of m^6^A_m_ modifications on RNA stability and translation (Drazkowska et al. 2022). The lack of effect on mRNA stability (Sendinc et al. 2019; Akichika et al. 2019; Sun et al. 2019) or the reduction of stability (Boulias et al. 2019) and translation was also shown by studies in HEK293, MEL624. Recent studies on PCIF1 knockout mice have shown that although m^6^A_m_ is not essential for the viability and fertility of mice, it does cause their reduced body weight. The authors explain the observed phenomenon by the RNA stabilizing role of m^6^A_m_ (Radha Raman Pandey et al. 2020) The lack of consensus in the accumulated data suggests that the recognition of the modified transcript may be subject to cell line-specific regulation (Doxtader & Nam, 2019; Tat & Kiss, 2021). Therefore, studies at the molecular level without the influence of other factors are required.

In this work, we undertook the investigation of the significance of cap1 and m^6^A_m_ modifications for the interaction between RNA and IFIT proteins. We studied interactions with both the IFIT1 protein alone as well as in complexes with IFIT2 and IFIT3. For this purpose, we used an assay we had previously developed that allowed us to perform the full kinetics of interactions between IFITs and RNA, as described in (Miedziak et al. 2019). The results obtained showed that the CA-m^6^A_m_ modification is a stronger signature than cap1 alone. It blocks the formation of a complex between IFIT proteins and m^7^Gpppm^6^A_m_-RNA several to dozens times more effectively than other studied by us cap modifications.

The model structure of the IFIT1-m7GpppApN complex (Abbas et al. 2017) shows water-mediated hydrogen bond formation between the N6 (A+1) position and V372. According to our conjecture and our results, the introduction of a methyl group into the N6 position of adenosine significantly weakens the formation of hydrogen bonds and, since the aforementioned hydrogen bonds were formed in the presence of a mediator such as water, the formation of additional hydrophobic interactions is impossible. Moreover, modelling with cap1-mRNA demonstrated that the methyl group would interfere with the amino acid residues R187 and Y157, introducing a steric hindrance and disrupting the interaction network through water molecules. Therefore, when the double methyl modification in the form of m^6^A_m_ is introduced into RNA, the formation of a stable IFIT1-RNA complex is no longer possible, which explains the strong decrease in the interactions observed in our study.

We also tested how the presence of modified m^6^A bases in the mRNA body (positions: +4, +6, +16) affects recognition by IFITs. We observed only a slight change in the kinetic parameters of m^7^GpppAG_m^6^A-RNA as compared to unmodified m^7^GpppAG-RNA. Among the other modified A, only +4 was located within the terminal part of the cap-binding channel, the others were already outside the channel. The slight modification of interaction parameters is most likely due to m6 presence within adenosine +4, resulting in rearrangement of the interaction network inside the channel. Especially since IFIT1’s crystallographic structure revealed the presence of adenine-specific hydrogen bonds between protein and N4 position of RNA. The 5ʹ untranslated region (UTR) is essential for translation initiation efficiency. The m^6^A-seq studies have revealed m^6^A peaks in the 5ʹ UTR and start codons (Zhou et al. 2015). Interestingly, m^6^A in the 5’ UTR could facilitate cap-independent translation through a process involving eIF3 (Meyer et al. 2015). In contrast, another report showed that the presence of a single m^6^A modification in the 5’ UTR did not affect translation dynamics (Guca et al. 2024). Our results suggest that the effect of m^6^A on translation is not due to this modification in the 5’ UTR acting through translation repression by IFIT proteins. The presence or absence of m^6^A in the 5’ UTR is not detected by IFIT proteins and this modification does not contribute to translation repression by IFIT proteins.

It has been proposed that IFIT3 binding to IFIT1 results in a specific increase in the affinity of the complex for m^7^GpppG-RNA, preferring this construct over others (e.g., cap1-RNA (Johnson et al. 2018)). The data collected by us indicate that the presence of IFIT3 increases the affinity of the complex towards all capped RNAs tested. None of the studied RNAs was favored over the other ones by the IFIT1/IFIT3 complex. Instead, the presence of IFIT3 was found to affect the dissociation of the protein-RNA complex. Specifically, IFIT3 slowed the off-rate of IFIT1 dissociation from RNA. This can be explained by the fact, that IFIT3 induces a tighter conformation of the IFIT1 protein on RNA.

In summary, we have characterized the interactions between the proteins of the IFIT complex, and determined the correct conditions for the formation of these complexes. We then determined how IFIT2 and IFIT3 proteins affect the specificity of IFIT1’s recognition of the 5’ end of RNA. One of the most interesting results is the comparison of the recognition of RNA with CA-m^6^A_m_ and cap1 by IFIT proteins. Our measurements show that CA-m^6^A_m_ is clearly a stronger signature than cap1 alone and is sufficient for RNA to remain unrecognized by IFIT1 or its complexes. Finally we showed that presence or absence of m^6^A in 5’ UTR is not detected by IFIT proteins and this modification does not contribute to translation repression by IFIT proteins.

## Materials and methods

### Solvents and Chemical reagents

Chemical reagents, including fully protected 2’-O-methyladenosine phosphoramidite were purchased from commercial sources. P-Imidazolide of N7-methylguanosine 5′-diphosphate (im-m^7^GDP), fully protected of 2′-*O,N^6^*-dimethyladenosine phosphoramidite and N2-isobutyryl-2′,3′-isopropylidene-guanosine were synthesized according to the previously described protocols (Kowalska et al. 2008; Eisenführ et al. 2003; Sikorski et al. 2020).

### General procedure of pNpG synthesis

The synthesis of dinucleotides was carried out in a solution using phosphoramidite chemistry. In the coupling step, 1.0 equivalents of 5′-O-DMT-2′-*O*-methyl/*N*6,2′-*O*-dimethyladenosine-3′-O-phosphoramide, 1.0 equivalents of protected guanosine and 0.40 M 5-(benzylthio)-1-H-tetrazolium in acetonitrile were mixed at room temperature under an argon atmosphere. After 4 h, the mixture was cooled to 4 °C and 0.1 M iodine in pyridine was added and stirred for 1h at RT. The reaction mixture was extracted with dichloromethane and washed with brine solution. The resulting organic layer was dried, evaporated and purified by flash chromatography on silica gel using gradient elution (0-5 % methanol in dichloromethane). The resulting solid was dissolved in a 20 % aqueous solution of TFA and stirred at RT for 4 h. The mixture was evaporated under reduced pressure and co-evaporated 6 times with methanol. The crude dinucleotide was dissolved in methanol and dropped into diethyl ether. The precipitate was filtered, washed with diethyl ether and dried in a vacuum desiccator over phosphorus pentoxide. In the final step, the dinucleotide was phosphorylated at the 5’-OH position by standard Yoshikawa method according to previously described protocols (Grzela et al. 2022). The resulting product was deprotected with ammonia, evaporated and purified by ion exchange chromatography on DEAE-Sephadex (A-25, HCO^3-^ form) using a linear gradient of triethylammonium bicarbonate (TEAB), pH 7.5, in water. Fractions containing the desired product were combined, evaporated and lyophilized to obtain TEA salt of the product as a fine white powder.

### General synthesis of trinucleotide cap analogues

Triethylammonium salt of pNpG (1.0 eq.), im-m^7^GDP (2.0 eq.) and anhydrous ZnCl_2_ (25 eq.) was dissolved in anhydrous DMF. The mixture was stirred at RT for 24 h and then the reaction was quenched by addition of aqueous solution of EDTA. The product was isolated by ion-exchange chromatography on DEAE Sephadex (gradient elution 0–1.2 M TEAB) and purified by semi-preparative RP HPLC (SUPELCOSIL™ LC-18-DB, gradient elution 0–50 % methanol in 0.05 M ammonium acetate buffer pH 5.9) to render, after evaporation and repeated freeze-drying from water, ammonium salt of trinucleotide cap analogues. The reaction yields ranged from 40 to 48 %.

### Protein expression and purification

pET28a(+)6xHis-TEV-IFIT1, pET28a(+)6xHis-TEV-IFIT2, and pET28a(+)6xHis-TEV-IFIT3 were a gift from Kathleen Collins (Addgene plasmid # 53557; http://n2t.net/addgene:53557; RRID:Addgene_53557, Addgene plasmid # 53558; http://n2t.net/addgene:53558; RRID:Addgene_53558; Addgene plasmid # 53559; http://n2t.net/addgene:53559; RRID:Addgene_53559)(Katibah et al. 2013). IFITs were expressed in Rosetta 2 (DE3)pLysS *Escherichia coli* (Novagen). Cells were grown to an OD_600_ of approximately 0.6 in LB media at 37 °C. Expression was induced by adding 0.2 mM isopropyl-D-1-thiogalactopyranoside. The induced culture was incubated at 16 °C for 16 h. Cells were harvested and lysed in buffer containing 50 mM sodium phosphate pH 7.5, 400 mM NaCl, 5 % glycerol, 0.5 mM DTT, 1 tablet of EDTA-free protease inhibitor, 6 µL Viscolase, 0.5 % Triton-X100 and a little bit of lysozyme. Next, cells were lysed by sonication with 60 % power for 5 x 90 seconds on ice, followed by centrifugation at 13 000 rpm for 30 minutes at 4 °C. The supernatant was collected, filtered and subsequently purified using His-trap column or Ni-NTA Agarose beads equilibrated with buffer (50 mM sodium phosphate pH 7.5, 400 mM NaCl, 5 % glycerol and 0.5 mM DTT), washed extensively with buffer (50 mM Sodium phosphate pH 7.5, 1 M NaCl, 10 mM Imidazole, 5% glycerol, 0.5 mM DTT) and finally eluted with buffer (50 mM sodium phosphate pH 7.5, 400 mM NaCl, 600 mM Imidazole, 5% glycerol and 0.5 mM DTT). Then the buffer was exchanged to 50 mM sodium phosphate pH 7.5, 150 mM NaCl, 0.5 mM DTT and 10 % glycerol on PD-10 desalting columns. To obtain IFITs ligands without his tag used for MST experiment, IFIT1 and IFIT2 were further cleaved with SUMO and TEV proteases, respectively. Each protein was additionally purified by size exclusion chromatography (SEC) on Superdex 200 increase 10/300 GL in SEC buffer (50 mM sodium phosphate pH 7.5, 150 mM NaCl, 0.5 mM DTT and 10 % glycerol) and concentrated on Amicon Ultra 15 mL Centrifugal Filters (Merck Millipore) with Nominal Molecular Weight Limit of 10 kDa according to the instruction. After purification, the presence and purity of desired IFIT proteins were confirmed by SDS polyacrylamide gel electrophoresis (SDS-PAGE).

### RNA preparation

Linear pSPluc+ plasmid template containing T7 class III promoter φ6.5 was obtained by restriction with XhoI and EcoRI. m^7^GpppAG-, m^7^GpppA_m_G-, m^7^Gpppm^6^A_m_G-capped mRNAs and m^7^GpppAG-caped RNAs with N^6^-methyladenosine in the RNA body were synthesized by *in vitro* transcription (IVT) with co-transcriptional capping, using 0.5 mM m^7^GpppAG, m^7^GpppA_m_G or m7Gppp^m6^A_m_G, 0.1 mM GTP, 12.5ng/µL linear pSPluc+ plasmid template, 0.5 mM ATP, CTP, UTP, ^m6^ATP, 1 U/µL T7 RNA Polymerase (Thermo), and 1U/µL RiboLock RNase Inhibitor (Thermo). After transcription, all RNAs were treated with alkaline phosphatase to remove any remaining phosphate groups from RNA. Finally, RNAs were purified using Oligo Clean-up and Concentration Kit (Norgen Biotek) and analyzed by denaturing PAGE on a 15% polyacrylamide/7 M urea gel. The resulting sequence of RNA is the following: (G/A)GGAGACCGGCCTCGA.

### Microscale thermophoresis (MST)

MST test was used to study protein interactions of IFITs. Variants of IFIT proteins used for MST test included his-tagged proteins (his-IFIT2 and his-IFIT3) and proteins without his tag as ligands (IFIT1 and IFIT2). Prior to the preparation of IFIT complexes, all proteins were centrifuged for 10 min at 12000 rpm and 4 °C to remove aggregates and precipitates. 200 µL of 100 nM solution of his-tagged protein was labeled with RED-tris-NTA 2^nd^ Generation dye (Monolith HIS-tag labeling Kit, NanoTemper). Protein-protein complexes were obtained by mixing dye-labeled IFIT protein (final concentration 50 nM) with 16 serial dilutions of another IFIT protein without a tag (ligand protein). The results obtained were used to determine the binding parameters for various IFIT complexes. 10 µL of each ligand protein dilution was mixed with 10 µL labeled protein and incubated at different temperatures (4 °C, 15 °C, 37 °C, 30 °C) for 30 min or subjected to further procedures without incubation. For hetero-complex of IFIT1/2/3, his-tagged IFIT3 labeled with dye was firstly incubated with IFIT2 in 1:1 molar ratio at 4 °C or at 15 °C to form heterodimer complex, followed by centrifugation (4 °C, 12000 rpm, 10 min) and then incubated at 4 °C or 15 °C with serial dilutions of IFIT1 to obtain final hetero-trimer. All protein complexes were centrifuged for 10 min at 12000 rpm at 4 °C and the upper solution of samples was further loaded into capillaries for measurements. The following binding studies were performed with a Monolith NT.115 device (Nano temper Technologies) using Monolith™ NT.115 Premium Capillaries (NanoTemper). Results with one inflection point were analyzed with the standard NanoTemper software using K_D_ model assuming one binding site or equivalent binding sites. Since we observed two separated inflection points for some samples, their analysis with standard NanoTemper software was insufficient. Therefore, the results were also analyzed using the model of two independent binding sites, as described previously by (Winiewska et al. 2017). Analysis was performed using Origin package (OriginLab, Northampton, MA, www.originlab.com) and dissociation constants were fitted globally for a series of at least three MST pseudo-titration experiments for each complex.

### Differential scanning fluorescence (DSF)

The protein thermal stability studies for all IFIT samples and complexes were performed using Prometheus NT.48 nanoDSF device (NanoTemper Technologies) with simultaneous Dynamic light scattering (DLS) measurements. Prior to the preparation of samples for measurements, all proteins were centrifuged with 12000 rpm at 4 °C for 10 min to remove aggregates and precipitates. IFIT protein complexes were obtained with or without the pre-incubation step performed in the same manner as in MST experiments. Single IFIT protein samples and IFIT complexes diluted to 5 µM concentration in buffer (50 mM sodium phosphate pH 7.5, 150 mM NaCl, 5% glycerol) were prepared and loaded into standard capillaries (NanoTemper Technologies). During the measurements, the thermal unfolding of the protein was monitored in the temperature range of 20–95 °C at a rate of 1 °C/min, with the excitation power of 30 %. The data processing procedure was performed using NanoTemper software dedicated to nanoDSF equipment, PR.ThermControl v2.1.2 (NanoTemper Technologies) and melting point temperatures (*T_m_*) were calculated as the minimum of the first derivative plot using the Origin 2019 package (OriginLab, Northampton, MA; www.originlab.com).

### Biolayer interferometry (BLI) assay

#### Protein complexes preparation

All IFIT proteins used for BLI assay were his-tagged (his-IFIT1, his-IFIT2 and his-IFIT3). Prior to the preparation of IFIT complexes, all proteins were centrifuged for 10 min at 12000 rpm, at 4 °C to remove aggregates and precipitates. IFIT1/3 complexes in different concentrations (0– 1200 nM) were obtained by incubation of IFIT1 with IFIT3 in a molar ratio of 1:1 at 15 °C for 30 min. IFIT1/2/3 trimers were obtained through two-step incubation. Firstly, IFIT2 mixed with IFIT3 in a molar ratio of 1:1 was incubated at 15 °C for 30 min and then, IFIT1 was incubated with IFIT2/3 complexes at 15 °C for 30 min to form the final trimer IFIT1/2/3 (1:1:1 molar ratio). All the complexes were centrifuged for 10 min at 12000 rpm, at 4 °C prior to the measurements.

#### Interaction assay

High Precision Streptavidin (SAX) Biosensors (Pall ForteBio) and the Advanced Kinetics module were applied on BLItz platform (Pall ForteBio) for BLI interaction assays. Short RNAs produced by IVT were biotinylated using Pierce RNA 3′ End Biotinylation Kit following the manufacturer’s guidance (Thermo). Immobilization of biotin-labeled RNAs containing different 5′ ends onto streptavidin-coated biosensors (Pall ForteBio) was achieved by immersing the sensors for 5 min with shaking at 1000 rpm in the kinetic buffer (50 mM phosphate buffer pH 7.2, 150 mM NaCl, 10 % glycerol, 0.5 mM DTT, 0.1 % BSA, and 0.05 % Tween 20) containing 1 µM RNA. Nonspecific binding was reduced by blocking the sensors with 10 µg/mL EZ-LINK Biocytin (Thermo) and subsequent washing with kinetic buffer. The association between RNA and proteins (IFIT1 alone, IFIT1/3 and IFIT1/2/3 complexes) was measured by incubation of RNA immobilized on sensors with a series of protein dilutions in the kinetic buffer. Afterwards, the biosensors were incubated in the kinetic buffer for 5 min to measure the dissociation constants. Global fitting in the ForteBio analysis software was used to analyze the data and determine equilibrium dissociation constant *K*_D_, kinetic association *k*_a_ and dissociation rates *k*_d_. All presented values are the means from three independent experiments.

## Authorship contribution statement

**Jingping Geng**: Investigations, Formal analysis, Writing – review and editing. **Magdalena Chrabaszczewska**: Investigations, Formal analysis, Methodology, Writing – original draft, Writing – review and editing. **Karol Kurpiejewski**: Investigations, Writing – review and editing. **Marzena Jankowska-Anyszka**: Conceptualization, Methodology, Writing – review and editing, Funding acquisition. **Anna Stankiewicz-Drogon**: Investigation, Formal analysis, Writing – review and editing. **Edward Darzynkiewicz**: Methodology, Writing – review and editing, Funding acquisition. **Renata Grzela**: Investigation, Formal analysis, Conceptualization, Data curation, Methodology, Project administration, Supervision, Writing - original draft, Writing – review and editing

## DATA AVAILABILITY

Data will be made available on request

## Acknowledgment

This work is dedicated to Edward Darzynkiewicz who passed away during the writing of this manuscript. Edward has dedicated his life to the study of cap structures and has made significant contributions to the understanding of the diverse processes in which RNA molecules are involved. pET28a(+)6xHis-TEV-IFIT1, pET28a(+)6xHis-TEV-IFIT2, pET28a(+)6xHis-TEV-IFIT3 were a gift from Kathleen Collins (University of California) (Addgene plasmids #53557, #53558, #53559). We thank Dr. Zbigniew Darzynkiewicz for reading the manuscript.

## Abbreviations

BLI: biolayer interferometry
CA-m^6^A_m_: Cap adjacent-m^6^A_m_
Cap1: m^7^GpppN_m_N
Cap2: m^7^GpppN_m_N_m_
CAPAM: cap-specific adenosine methyltransferase
CMTR1: cap 2’-*O*-methyltransferase 1
DLS: dynamic light scattering
DMF: N,N-dimethylformamide
DSF: differential scanning fluorimetry
DTT: 1,4-dithiothreitol
EDTA: ethylenediaminetetraacetic acid
FTO: Alpha-ketoglutarate-dependent dioxygenase (fat mass and obesity-associated protein)
IFIT: interferon-induced proteins with tetratricopeptide repeats
ISG: IFN-stimulated genes
ITC: isothermal titration calorimetry
IVT: in vitro transcription
*k*_a_: association rate constant
*k*_d_: dissociation rate constant
*K*_D_: equilibrium dissociation constant
m^6^A: *N*6-methyladenosine
m^6^A_m_: *N*6,2’-*O*-dimethyladenosine
MST: microscale thermophoresis
NTA: nitrilotriacetic acid
PAMP: Pathogen-associated molecular patterns
PCIF: phosphorylated CTD-interacting factor 1
PRR: pattern recognition receptor
RIG-I: retinoic acid-inducible gene I
RP-HPLC: reverse phase high-performance liquid chromatography
RPM: revolutions per minute
RT: room temperature
SAX: high precision streptavidin
SDS-PAGE: SDS polyacrylamide gel electrophoresis
SEC: size exclusion chromatography
SUMO: small ubiquitin-like modifier
TEA: triethylamine
TEAB: triethylammonium bicarbonate
TEV protease: Tobacco Etch Virus nuclear-inclusion-a endopeptidase
TFA: trifluoroacetic acid
*T*_agg_: aggregation temperature
*T*_m_: melting point temperature

